# Integrated Characterization of Proximal and Distal Nerve Stumps in a Model of Chronic Motor Nerve Transection

**DOI:** 10.64898/2026.11.02.763804

**Authors:** Chieh Yu, Lenae Joe, Anthony Newman, Jon-Paul Pepper

**Affiliations:** Department of Otolaryngology - Head & Neck Surgery, Stanford University School of Medicine, Palo Alto, CA, 94304 United States

**Keywords:** Chronic nerve injury, Axotomy, Denervation, Neurotmesis, Motor axons, Myelination, Traumatic neuroma, Endoneurial remodeling, Extracellular matrix, Axon morphometry, Neuropathology, Preclinical model

## Abstract

Prolonged peripheral nerve discontinuity subjects the proximal stump to chronic axotomy and the distal stump to chronic denervation, resulting in pathological changes that may compromise regeneration following delayed repair. Although these changes have been investigated in both nerve stumps, much of the existing evidence derives from mixed sensorimotor nerve models, leaving the histopathological changes of chronic motor nerve injury incompletely understood. Addressing this gap requires a well-defined, reproducible model that maintains prolonged motor nerve discontinuity and enables concurrent characterization of both nerve stumps. This study aimed to establish a chronic motor nerve transection model and characterize the distinct structural and microenvironmental changes occurring in both nerve stumps. The buccal branch of the facial nerve was transected in Lewis rats, and the nerve ends were secured to adjacent muscle to prevent spontaneous reconnection. At 10 weeks post-injury (p.i.), proximal and distal stumps were compared with uninjured nerves using histological, immunofluorescence, and morphometric analyses. Persistent functional impairment was observed throughout the study period. The proximal stump exhibited traumatic neuroma formation, altered axonal morphology, and loss of myelin integrity, but largely preserved its underlying nerve architecture and microenvironment. In contrast, the distal stump exhibited reduced neurofilament expression, clusters of small-caliber myelinated axons, and increased myelinated axon density compared to both uninjured nerves and the proximal chronically injured segment. These changes were accompanied by endoneurial tube atrophy, increased collagen I deposition, vascular remodeling, and immune-cell accumulation, indicating extensive structural and microenvironmental remodeling. Collectively, these findings demonstrate that chronic motor nerve transection produces two distinct pathological compartments: a proximal stump characterized by persistent but disorganized regeneration-associated features and a distal stump characterized by pronounced degenerative and microenvironmental changes. This study establishes a reproducible model of chronic peripheral nerve injury and provides a foundation for investigating how these compartment-specific pathologies contribute to regenerative failure and for developing targeted therapeutic strategies to improve regeneration following delayed nerve repair.

**Graphical Abstract:** Distinct Proximal and Distal Stump Pathology in a 10-Week Chronic Motor Nerve Transection Model

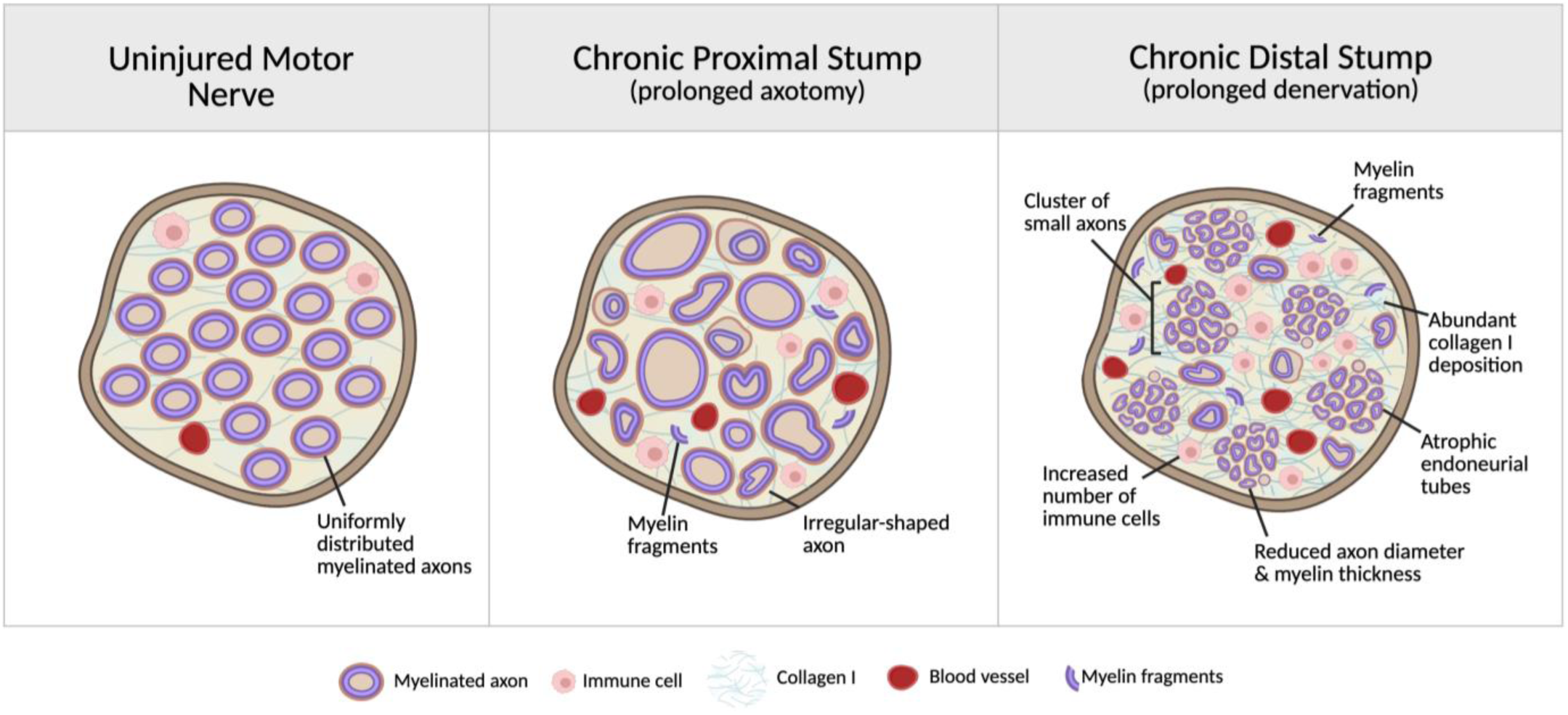

## Introduction

Severe peripheral nerve injuries (PNIs), particularly those involving nerve transection or segmental defects, are associated with a greater risk of persistent functional deficits, especially when timely surgical repair is not feasible (Aldali et al., 2026; Rochkind, 2026; Yang et al., 2026). Delayed nerve repair remains a significant clinical challenge, as prolonged nerve discontinuity is associated with progressive changes that diminish regenerative outcomes (Sulaiman and Gordon, 2000; Gordon et al., 2011; Jonsson et al., 2013; Lynch and Mahmood, 2026). In motor nerve injuries, functional recovery depends on the ability of regenerating axons to traverse the distal nerve stump and reinnervate their target muscles. However, prolonged nerve discontinuity exposes the proximal and distal stumps to two distinct but interrelated pathological processes: chronic axotomy and chronic denervation, respectively. Chronic proximal axotomy can lead to persistent axonal sprouting, structural disorganization, and traumatic neuroma formation, whereas prolonged distal denervation is associated with declining Schwann cell regenerative support, extracellular matrix remodeling, and fibrosis (Terenghi et al., 1998; Ronchi and Raimondo, 2017; Deininger et al., 2024). These divergent responses highlight the two-compartment nature of chronic PNI, in which pathological changes in either stump may compromise nerve regeneration. Characterization of both proximal and distal nerve stumps is therefore essential for a more complete understanding of the pathological changes associated with chronic nerve injury.

Existing experimental studies have provided valuable insights into chronic PNI, with much of this work emphasizing distal stump degeneration and the decline in regenerative support during prolonged denervation. Complementing this distal-focused literature, paired analyses have demonstrated distinct proximal and distal responses, particularly during acute and subacute injury stages. For example, Aiki *et al*. (2018) identified site-and stage-specific protein expression patterns in both stumps at 5, 10, and 35 days after sciatic nerve transection, while Cheng *et al*. (2021) compared cytokine expression between the two compartments over 1–28 days after injury. Chronic comparative studies have further demonstrated compartment-specific pathology. Terenghi *et al*. (1998) examined Schwann cells and axonal regeneration in proximal and distal stumps from human nerves undergoing reconstruction 8–53 months after injury, while Jonsson *et al*. (2013) assessed both compartments in a rat delayed sciatic nerve repair model. However, much of this experimental work has used mixed nerves containing both sensory and motor axons, which can complicate interpretation of changes specifically associated with motor nerve injury (Hercher et al., 2020; Li et al., 2025; Yang et al., 2025). A motor-specific model offers a more defined platform for examining chronic pathology relevant to motor axon regeneration and muscle reinnervation, yet integrated characterization of proximal and distal structural and microenvironmental changes in such models remains limited. Addressing this gap requires a reproducible model of chronic unrepaired motor nerve injury that maintains prolonged discontinuity and enables direct comparison of both compartments.

To meet this experimental need and address the biological gap, we established a rat model of chronic motor nerve transection using the buccal branch of the facial nerve, a surgically accessible motor nerve. Our objectives were to determine whether the model reproduces recognized pathological hallmarks of prolonged nerve discontinuity and to systematically characterize the compartment-specific structural and microenvironmental changes associated with chronic axotomy and denervation. At 10 weeks post-injury (p.i.), a stage of chronic denervation reported in previous studies (Wagstaff et al., 2021; Xu et al., 2023; Duan et al., 2026), the model exhibited persistent functional impairment and distinct proximal and distal pathological states. Proximal stumps showed relative preservation of nerve architecture alongside neuroma-associated remodeling, whereas distal stumps exhibited pronounced axonal, endoneurial, extracellular matrix, vascular, and immune-associated changes. Together, these findings establish a motor-focused experimental platform that integrates sustained nerve discontinuity with paired proximal–distal characterization, providing a foundation for investigating chronic injury progression and developing delayed-repair and compartment-targeted therapeutic strategies.

## Methods

### Experimental Animals

All animal procedures were approved by the Stanford University Administrative Panel on Laboratory Animal Care (APLAC; Protocol No. 32953) and were conducted in accordance with institutional guidelines for the care and use of laboratory animals. Male and female Lewis rats (9–10-weeks old) were purchased from Charles River Laboratories (Wilmington, MA, USA). Animals were housed under standard laboratory conditions with a 12-hour light/dark cycle and provided standard rodent chow and water *ad libitum*. The rats were acclimated for at least one week before surgery.

### Facial nerve transection injury surgery

All surgical procedures were performed under standard aseptic conditions. At the time of surgery, female rats weighed 230–250 g, and male rats weighed 360–420 g. Rats were anesthetized with isoflurane and sustained-release buprenorphine (0.5 mg/kg body weight) was administered subcutaneously for analgesia. Fur over the buccal region was shaved, and a mid-buccal skin incision was made to expose the buccal branch of the facial nerve. The buccal branch was completely transected, and the proximal and distal nerve stumps were diverted in opposite directions and secured to adjacent muscle beds using 9-0 nylon sutures (Synergy Surgical, Cat.# AA-0116) to prevent spontaneous reinnervation (Fig. 1). The skin incision was closed with 6-0 polyglactin (PGLA) sutures (Synergy Surgical, Cat. # G6031), and the animals were allowed to recover. Animals were euthanized by CO₂ inhalation at 10-weeks p.i. for the chronic time point (Fig. 1A), followed by bilateral pneumothorax to confirm death. The 10-weeks p.i. timepoint was selected based on previous studies demonstrating that regenerative impairment is evident by this stage of chronic denervation, including reduced expression of the repair-associated c-Jun in Schwann cells (Wagstaff et al., 2021; Xu et al., 2023; Duan et al., 2026).

**Figure 1.**
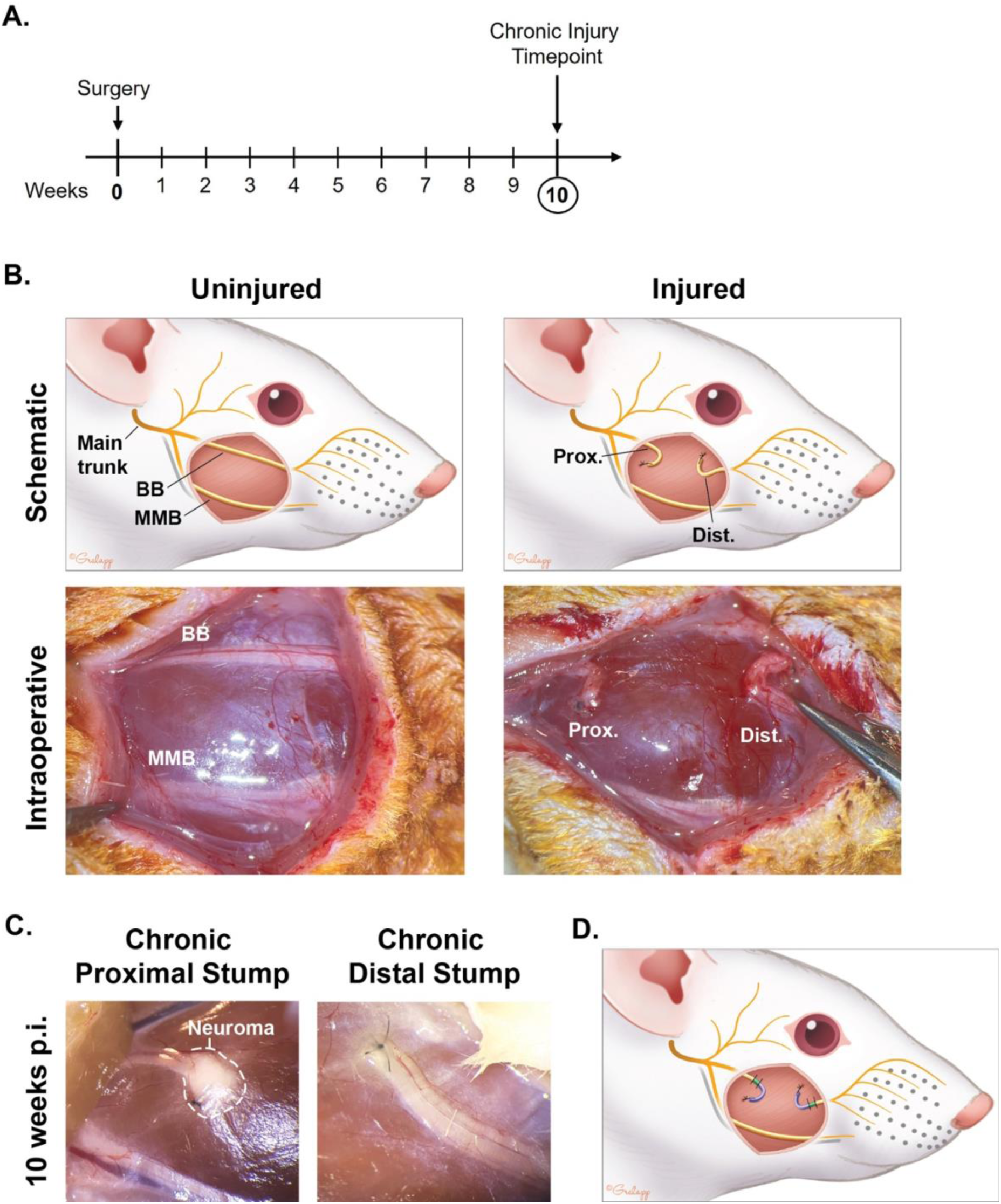
Establishment of a rat facial nerve chronic transection injury model. **(A)** Animal surgical timeline. **(B)** Schematic and representative intraoperative images of the uninjured and injured buccal branches. **(C)** Representative gross images of the proximal and distal buccal nerve stumps at 10-weeks p.i. **(D)** Schematic of tissue sampling from the proximal and distal buccal branch stumps. Segments indicated in purple were used for longitudinal sectioning, while segments indicated in green were used for cross-sectional analysis. BB, buccal branch; MMB, marginal mandibular branch; p.i., post-injury. Rat illustrations (B, D) published with permission from Ms. Christine Gralapp.

### Whisker function analysis

For functional assessment purpose, in addition to the buccal branch transection, the marginal mandibular branch was transected and suture repaired to completely ablate nerve input to the vibrissae muscle (Supplementary Fig. 1). To assess functional recovery, longitudinal video recordings of rat whisking were obtained weekly from 1- to 10-weeks p.i. using an iPhone camera in slow-motion mode (Supplementary Video 1). Rats were placed in a black box and recorded using the Mono filter to improve whisker contrast. Rats were allowed to walk around freely, with chow used to entice whisking. Video recordings were reviewed and manually scored in a blinded manner. Rats were scored on a 1-5 scale for movement and position of the affected vibrissae as previously described (Faria et al., 2006). Briefly, a score of 1 for movement corresponded to no vibrissae movement, whereas a score of 5 corresponded to complete and symmetric vibrissae movement. The intermediate scores, 2 through 4, were assigned for relative movement of the affected vibrissae compared to the unaffected vibrissae. A score of 2 had slight shivering, a score of 3 had intermediate movement, and a score of 4 had almost symmetric movement. For position, vibrissae in a completely posterior position were scored a 1 and vibrissae in the anterior position and completely symmetric with the unaffected side were scored a 5. Scores 2, 3, and 4 were used to describe the relative position with greater specificity. A cohort of ten rats (n = 5 males and n = 5 females) were used for whisker function assessment.

### Hematoxylin and Eosin (H&E) staining

Facial nerve tissue was harvested and immediately fixed in 4% paraformaldehyde (PFA) at 4°C overnight. Tissue processing and histological staining was performed by Histo-tec Lab (Hayward, CA). After fixation, tissues were dehydrated through a graded ethanol series, cleared in xylene, and embedded in paraffin. Serial sections (5–10 µm thick) were cut using a rotary microtome and mounted on glass slides for histological staining. Paraffin-embedded tissue sections were stained with hematoxylin and eosin (H&E). Briefly, sections were deparaffinized, rehydrated through a graded ethanol series, stained with Richard Allan hematoxylin and Richard Allan eosin with phloxine, dehydrated, cleared in xylene, and coverslipped. Stained sections were examined under a brightfield microscope, and brightfield images were captured using AxioVision software (Carl Zeiss Micoscopy, LLC).

### Immunofluorescence Analysis

Longitudinal (15 µm) and transverse (10 µm) cryosections were incubated in blocking buffer (5% normal goat serum [NGS] and 1% bovine serum albumin [BSA] in 0.1% Triton X-100 in PBS [PBST]) for 1 h at room temperature (RT). Tissue sections were incubated overnight at 4°C with primary antibodies diluted in blocking buffer. Tissues were then washed three times for 10 min each in 0.1% Tween 20 in PBS and incubated for 1 h at RT with the appropriate secondary antibodies and Hoechst 33342 (1:1,000; BD Biosciences, Cat.# 561908, RRID:AB_2869394). After three washes for 10 min each with PBS, samples were mounted using Fluoromount-G mounting medium (SouthernBiotech, Cat.# 0100-01) and size no. 1.5 coverslips (VWR, Cat.# 48393-195) Tissue sections were imaged using an LSM 900 confocal laser scanning microscope (Carl Zeiss Microscopy GmbH, Jena, Germany). For each immunofluorescence marker, tissues from two male and two female rats (n = 4 biological replicates) were imaged using identical microscope settings and subsequently quantified.

Primary antibodies were chicken anti-neurofilament heavy (NFH) (1:1,000; Abcam, Cat# ab4680; RRID:AB_304560), rabbit anti-stathmin-2 (STMN2/SCG10) (1:300; Novus Biologicals, Cat.# NBP1-49461; RRID:AB_10011569), rabbit anti-laminin (1:300; Abcam, Cat.# ab11575; RRID:AB_298179), rabbit anti-collagen I (1:300; Abcam, Cat.# ab316222; RRID:AB_3697422), rabbit anti-CD31/PECAM-1 (1:300; Novus Biologicals, Cat.# NB100-2284; RRID:AB_10002513), and Alexa Fluor® 488-conjugated rabbit anti-Iba-1 (1:100; Abcam, Cat.# ab225260; RRID:AB_3750983). Secondary antibodies were Alexa Fluor® 594-conjugated goat anti-rabbit IgG (H+L) (1:1,000; Jackson ImmunoResearch Labs, Cat.#: 111-585-003; RRID:AB_2338059), and Alexa Fluor® 647-conjugated goat anti-chicken IgG (H+L) (1:1,000; Abcam, Cat.# ab150175; RRID:AB_2732800).

### Quantitative Image Analysis

Quantitative image analysis was performed using FIJI (ImageJ version 1.54p). Regions of interest (ROIs) were randomly selected for analysis. For longitudinal nerve sections, immunofluorescence signal area fraction (area %) was quantified for NFH, SCG10, and Iba-1 as the percentage of the total region of interest (ROI) area occupied by marker-positive staining. Three 100-µm-long ROIs were analyzed per biological replicate, with four biological replicates (n = 2 males and n = 2 females) examined for each marker. For transverse sections, collagen I and PECAM-1 signal area fraction was quantified from three ROIs per biological replicate across four biological replicates. A fixed threshold was established independently for each marker and applied consistently to all images within the corresponding analysis. For laminin-stained transverse sections, endoneurial tube area was measured within the inner boundary of the laminin-positive staining using three randomly selected ROIs per biological replicate across four biological replicates. Data were plotted using GraphPad Prism (v11.0.1, GraphPad Software, Boston, MA, USA). Measurements from individual ROIs were displayed as smaller data points and color-coded according to their corresponding biological replicate. The mean of the ROI measurements from each animal was used as the biological replicate value and displayed as a larger data point. Group means and standard deviations (SDs) were calculated from biological replicate values.

### Osmium tetroxide and Toluidine Blue O staining

Resin-embedded nerve tissue processing for semithin sections was performed by the Stanford University Cell Sciences Imaging Core Facility (Stanford, CA; RRID:SCR_017787). Briefly, dissected nerve tissues were fixed in 2% glutaraldehyde and 4% PFA in PBS, and post-fixed in 2% osmium tetroxide (OsO4) in cacodylate buffer for 1.5–1.75 h at room temperature. Samples were subsequently treated with 2.5% potassium ferricyanide in cacodylate buffer for 1.5 h, rinsed twice in distilled water, and incubated in freshly prepared 1% w/v thiocarbohydrazide for 20 min following dissolution at 60°C. After additional distilled water washes, tissues underwent a second osmium staining step in 2% OsO4 in water for 90 min at RT. En bloc staining was then performed with 1% aqueous uranyl acetate overnight at 4°C followed by incubation for 2 h at 50°C, washed in distilled water, and stained with Walton’s lead aspartate for 2 h at 50°C. Samples were dehydrated through a graded ethanol series (30%, 50%, 70%, 95%, and 2 × 100% ethanol) followed by acetonitrile. Tissue infiltration was performed using increasing ratios of Embed-812 hard resin to acetonitrile (1:3, 1:1, and 3:1), followed by 100% resin infiltration overnight. Samples were embedded in fresh Embed-812 resin and polymerized in molds. Resin blocks were trimmed and semithin sections (250nm) were counterstained with 1% toluidine blue O solution for 1 min at 70°C. Brightfield images were acquired using a Axio Imager upright light microscope equipped with AxioVision software (Carl Zeiss Microscopy GmbH, Jena, Germany) at 63× oil-immersion magnification.

### Morphometric analysis

For the segmentation and morphometric analysis of the nerve fibers, the AxonDeepSeg plugin (Zaimi et al., 2018) within the napari multi-dimensional image viewer (Sofroniew et al., 2026) was utilized. Automated segmentation was performed using the model_seg_generalist_BF_light pre-trained model. Following the initial AI-driven segmentation, axon and myelin masks were manually refined in napari to ensure anatomical accuracy; this included contour refinement and the elimination of false-positive detections. Digital measurements were converted to physical dimensions using image-specific pixel sizes calculated from the scale bar-derived physical dimensions of each region of interest and its corresponding pixel dimensions. For axons with irregular cross-sectional geometries, the axon diameter was mathematically derived from the total cross-sectional area of each individual axon (Zaimi et al., 2018). Fibers touching the ROI border or exhibiting incomplete profiles were excluded from morphometric analysis. A total of four ROIs across three biological replicates were analyzed for each sample condition. Final morphometric parameters were computed through the AxonDeepSeg integrated analysis tools, and all subsequent data visualization and statistical analyses were performed using GraphPad Prism 11 (v11.0.1, GraphPad Software, Boston, MA, USA).

### Statistical analysis and data presentation

Statistical analyses of immunofluorescence images quantification and morphometrics were performed using GraphPad Prism (version 11.0.1; GraphPad Software, Boston, MA, USA). Differences among uninjured, chronic proximal, and chronic distal nerve tissues were assessed using one-way group comparisons selected according to data distribution and variance assumptions. Technical replicates were averaged within each biological replicate, and biological-replicate values were used for all statistical analyses. Assumptions for parametric testing were assessed before group comparisons. Normality of model residuals was evaluated using the Shapiro–Wilk test and homogeneity of variance was assessed using the Brown–Forsythe test. Data meeting these assumptions were analyzed by ordinary one-way ANOVA with Tukey’s multiple comparisons test. When homogeneity of variance was violated, Brown–Forsythe and Welch ANOVA tests with Dunnett’s T3 multiple comparisons test were used. For whisker movement and position scores, a Friedman test was used to assess differences across time points, followed by Dunn’s multiple-comparisons test to compare each time point with 1-week p.i.. Data are presented as mean ± standard deviation (SD) unless otherwise indicated. Statistical significance was defined as \**p* < 0.05, \*\**p* < 0.01, \*\*\**p* < 0.001, and \*\*\*\**p* < 0.0001.

## Results

Chronic motor nerve transection creates prolonged nerve discontinuity, exposing the proximal stump to chronic axotomy and the distal stump to chronic denervation, two compartment-specific states that are each implicated in failed regeneration after delayed repair. Although prior delayed-repair and cross-anastomosis models have defined important independent contributions of proximal and distal degeneration, less is known about how these compartments appear side-by-side within a single unrepaired chronic motor nerve model, particularly beyond acute timepoints. To address this gap, we used a rat facial motor nerve transection model and characterized both proximal and distal stumps 10 weeks p.i. relative to uninjured nerve using complementary histological, immunofluorescent, and morphometric analyses to define compartment-specific axonal, cellular, structural, and microenvironmental tissue states relevant to prolonged discontinuity.

### Chronic facial nerve injury results in persistent denervation and neuroma formation

We established a chronic motor nerve injury model using the rat facial nerve buccal branch, with 10-weeks post-injury (10-weeks p.i.) defined as the chronic time point (Fig. 1A). Based on previous chronic denervation studies, regenerative impairment is evident by the 10-weeks p.i. timepoint (Wagstaff et al., 2021; Xu et al., 2023; Duan et al., 2026). The buccal branch was completely transected and the two nerve stumps diverted in opposite directions and sutured to nearby muscle bed to minimize re-innervation (Fig. 1B). By tethering the nerve stumps, it also allows for the preservation of nerve tissue for subsequent assessment or intervention. At 10-weeks p.i., proximal and distal buccal nerve stumps were collected and subjected to histological and immunohistological analyses. Grossly, the proximal stump exhibits a terminal traumatic neuroma, indicative of attempted regeneration (Huang et al., 2023) which is not present on the distal stump (Fig. 1C).

To functionally validate that transection and diversion of the buccal branch produces a persistent decrease in motor function, whisker movement and position were assessed weekly from 1- to 10-weeks p.i. (Supplementary Fig. 1A). Because the buccal and marginal mandibular branches both contribute motor input to the vibrissae musculature, buccal branch transection alone does not completely abolish whisker movement (Marin et al., 2026). Therefore, for functional assessment, the marginal mandibular branch was also transected and immediately suture-repaired to diminish its initial contribution to recovery, while allowing subsequent recovery through marginal mandibular branch regeneration (Supplementary Fig. 1B). This enabled persistent whisking deficits to reflect the continued absence of buccal branch input once marginal mandibular branch-mediated recovery occurred. However, a technical consideration is the close proximity of the marginal mandibular branch to the parotid duct, which is difficult to distinguish intraoperatively, but becomes more apparent during postmortem dissection. Consequently, manipulation of the marginal mandibular branch carries a risk of inadvertent duct injury and subsequent salivary leakage (Supplementary Fig. 1C).

Whisker movement and position were effectively abolished at 1-week p.i.. Functional recovery was significant by 3-weeks p.i., likely reflecting regeneration across the repaired marginal mandibular branch, but subsequently plateaued from 4- to 10-weeks p.i. with no further significant improvement (Supplementary Fig. 1D & 1E). At 10-weeks p.i., the mean whisker movement score was 2.64 ± 0.80 and the mean whisker position score was 3.09 ± 0.54, corresponding to 53% and 62% of normal function, respectively. An example of the persistent whisker function deficit is shown in Supplementary Video 1. Thus, although marginal mandibular branch repair permitted partial functional recovery, substantial whisker deficits persisted through 10-weeks p.i., consistent with continued discontinuity of the buccal branch.

Following confirmation of persistent functional deficit, histological and immunofluorescence analyses were performed on both proximal and distal nerve stumps. For tissue sampling, approximately 3–4 mm of each nerve stump immediately adjacent to the injury site was excised and processed for longitudinal sectioning (Fig. 1D, purple). Imaging was performed within the half of each excised segment farthest from the terminal cut end to better reflect the nerve segments that would typically be retained following nerve debridement during surgical repair (Fones et al., 2024). For transverse analysis, nerve segments beyond the excised 3–4 mm region were harvested and sectioned transversely (Fig. 1D, green).

### Differential axonal remodeling dynamics between chronic proximal and distal nerve stumps

General nerve morphology was assessed by hematoxylin and eosin (H&E) staining, revealing distinct architectural changes between uninjured nerves and chronically injured proximal and distal stumps (Fig. 2A). Uninjured nerves showed axons arranged in parallel, with relatively uniform cellularity and preserved nerve architecture. In chronic proximal stumps, nerve architecture appears relatively attached; however, the presence of digestion chambers indicated ongoing axonal and myelin degeneration and clearance. Degenerative changes were more pronounced in chronic distal stumps, which showed presence of digestion chambers, fragmented axonal profiles, and marked hypercellularity. Immunofluorescence analysis of neurofilament heavy chain (NFH), an axonal marker, revealed relatively preserved axonal architecture in the chronic proximal stump, whereas the chronic distal stump exhibited more fragmented and discontinuous NFH staining (Fig. 2B), indicating greater axonal degeneration distally. Quantification showed a lower NFH- positive area fraction in the chronic proximal stump (15.72 ± 5.09%) compared with uninjured nerves (22.03 ± 4.04%), although this difference was not statistically significant. In contrast, NFH-positive area fraction was significantly reduced in the chronic distal stump (9.50 ± 2.81%) fraction compared to uninjured nerves (*p* = 0.0048; Fig. 2C). To assess regenerative activity, SCG10, a marker associated with regenerating axons and extending growth cones (Shin et al., 2014), was examined. SCG10 staining was readily detectable in some chronic proximal stump samples, although expression varied across biological replicates, but was largely absent in the distal stump (Fig. 2D). Quantification of SCG10-positive area fraction showed no statistically significant difference between uninjured nerves (1.75 ± 1.61%) and both the chronic proximal stump (5.18 ± 3.53%) and the chronic distal stump (1.14 ± 1.69%) (Fig. 2E). Together, these findings highlight distinct pathological changes in the proximal and distal stumps following chronic nerve transection, with relatively preserved proximal nerve architecture but more pronounced axonal degeneration and structural disruption distally.

**Figure 2.**
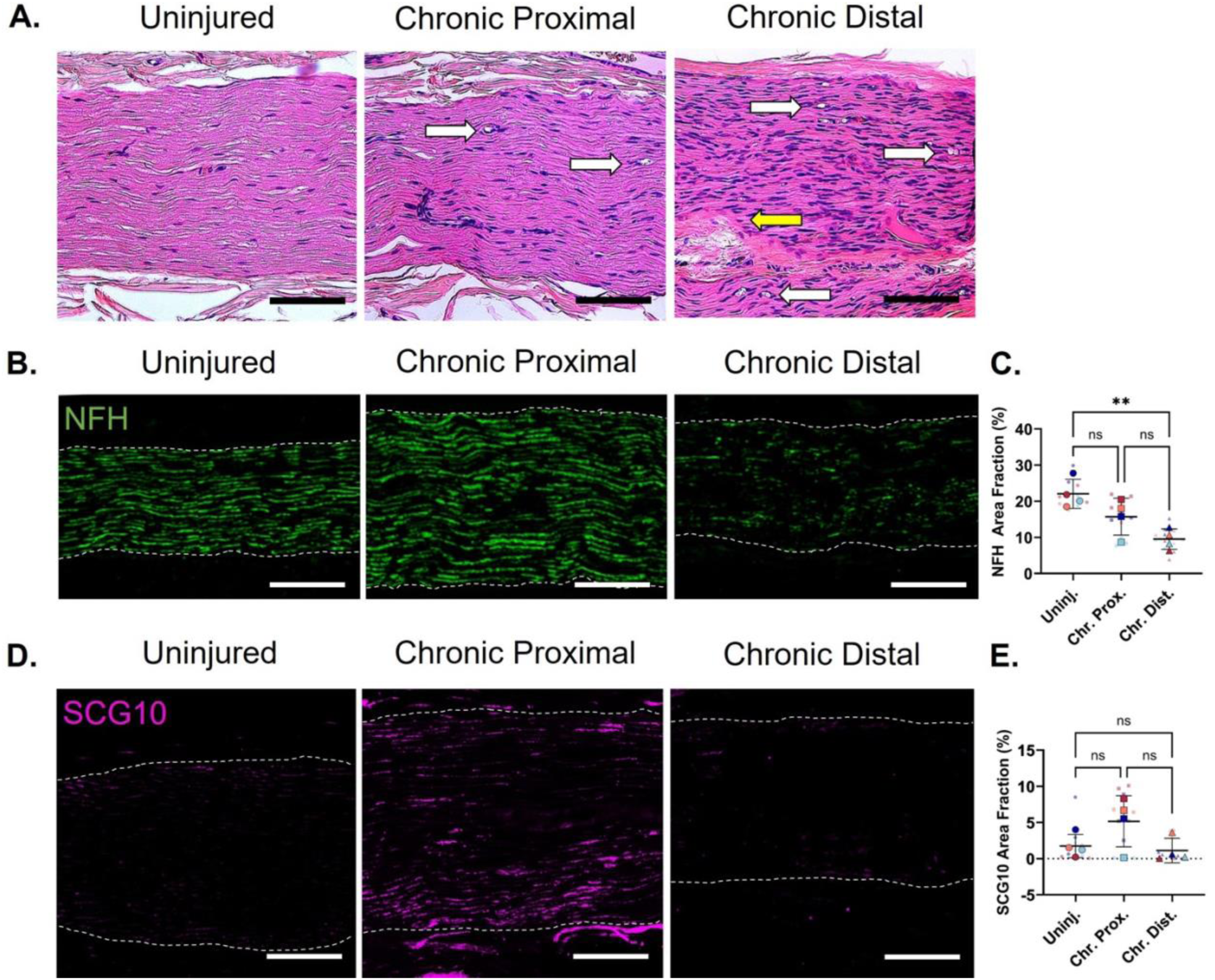
Distinct proximal and distal axonal remodeling states in the chronic transection model. **(A)** Representative brightfield images of longitudinal Uninjured, Chronic Proximal, and Chronic Distal nerve sections stained with hematoxylin and eosin (H&E). White arrows indicate myelin ovoids/digestion chambers, and the yellow arrow indicate axonal fragmentation. Representative immunofluorescence images and corresponding area-fraction quantification of **(B, C)** neurofilament heavy (NFH), highlighting axonal structure, and **(D, E)** stathmin-2 (STMN2/SCG10), highlighting regenerating axons, across Uninjured, Chronic Proximal, and Chronic Distal longitudinal nerve sections. White dashed lines indicate the boundaries of the nerve tissue. Scale bar = 100 µm. Data are presented as mean ± SD. Statistical significance is denoted as ** *p* < 0.01.

### Chronic nerve stumps exhibit distinct patterns of axonal and myelin remodeling

To further assess myelinated axon morphology and myelin integrity, we examined osmium tetroxide–postfixed, resin-embedded semithin nerve cross-sections stained with toluidine blue O. This analysis enabled detailed morphometric assessment and revealed distinct structural differences among uninjured nerves and chronic proximal and distal stumps. (Fig. 3A). Uninjured nerves exhibited relatively uniformly distributed myelinated axons of varying calibers, with predominantly rounded to oval axonal profiles surrounded by well-defined, continuous myelin sheaths. In contrast, chronic proximal stumps displayed greater morphological heterogeneity, characterized by irregularly shaped axonal profiles, and distorted and occasionally fragmented myelin sheaths. Chronic distal stumps exhibited more pronounced structural alterations, with numerous small-caliber myelinated axons frequently arranged in clusters, with less well-defined myelin sheath boundaries (Fig. 3A). To enable quantitative comparison of these morphological differences, toluidine blue images were segmented to identify individual axons (blue) and myelin profiles (red) for subsequent morphometric analysis (Fig. 3B). Axon density, expressed as the number of myelinated axons per mm^2^, was comparable between uninjured nerves (16,150 ± 3,493 axons/mm^2^) and chronic proximal stumps (15,967 ± 5,359 axons/mm^2^), but was significantly increased in the chronic distal stump (50,050 ± 7,375 axons/mm^2^) compared with both uninjured nerves and chronic proximal stumps (*p* = 0.0008 for both comparisons), consistent with the predominance of densely clustered small-caliber myelinated axons observed histologically (Fig. 3C).

**Figure 3.**
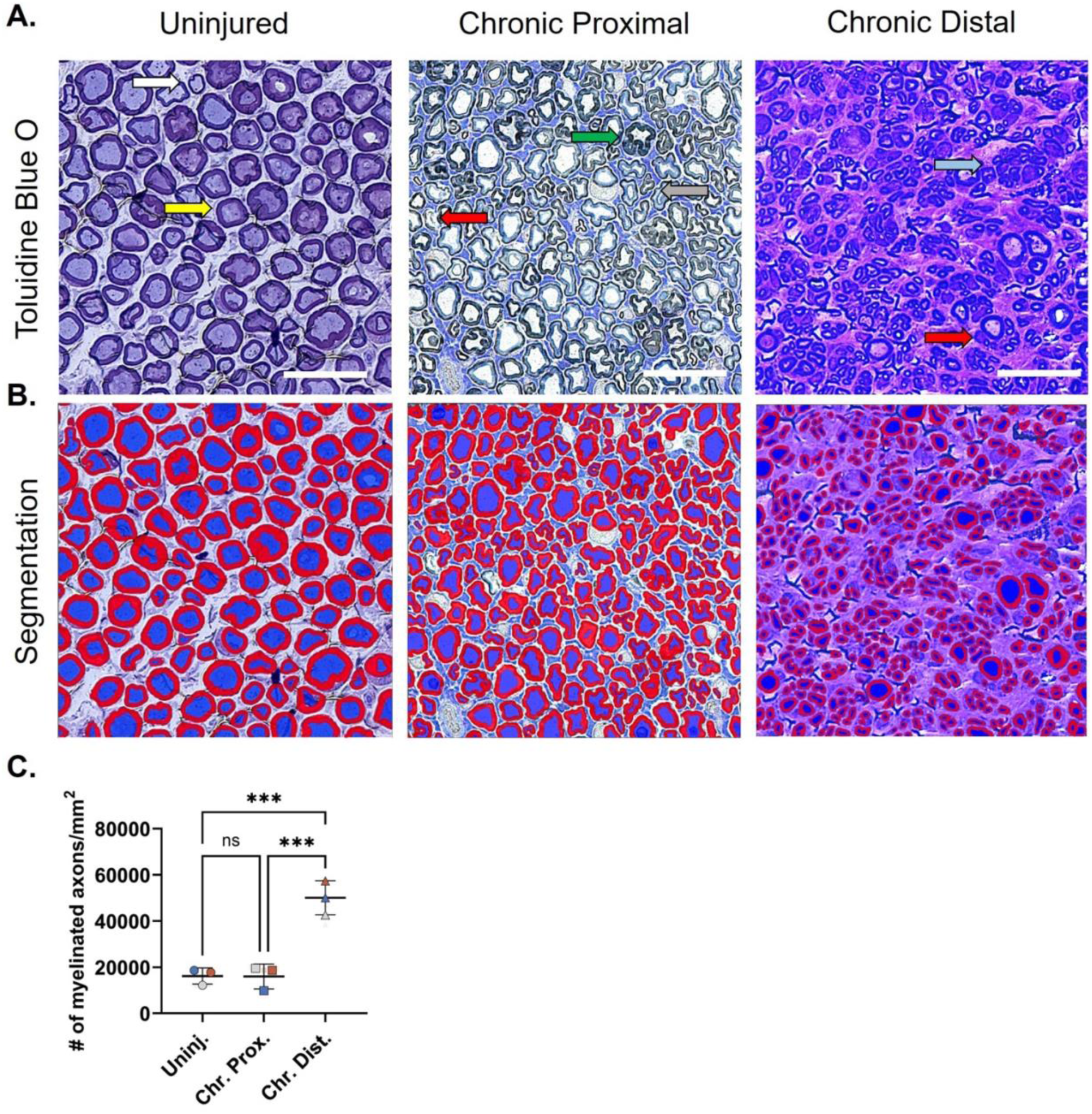
Compartment-specific changes in myelinated axon density and nerve architecture. **(A)** Representative toluidine blue O-stained images of transverse buccal nerve sections from Uninjured nerves, and Chronic Proximal and Chronic Distal nerve stumps. Yellow arrows indicate large-caliber axons with thick myelin; white arrows indicate medium-caliber axons with thinner myelin; green arrows indicate irregular myelin; gray arrows indicate small-caliber axons; red arrows indicate fragmented myelin; and light blue arrow indicate clusters of small-caliber axons. **(B)** Corresponding segmentation images generated using AxonDeepSeg, with axonal area shown in blue and myelin area shown in red. **(C)** Quantification of myelinated axon density (# of myelinated axons/mm^2^) in Uninjured, Chronic Proximal, and Chronic Distal nerve sections. Scale bar = 25 µm. Data is presented as mean ± SD. Statistical significance is denoted as *** *p* < 0.001.

Next, to quantify these morphological differences between the chronic proximal and distal stumps, we performed detailed morphometric analyses of myelinated axon caliber and myelin profiles. Analysis of axon diameter distributions further demonstrated differences in axon caliber among groups (Fig. 4A). Uninjured nerves exhibited a unimodal distribution peaking at 4 µm. The chronic proximal stump also exhibited a unimodal distribution, but with a shift toward smaller axon diameters, peaking at 3 µm. In contrast, the chronic distal stump displayed a pronounced leftward shift toward smaller axon diameters, with approximately 70% of axons having diameters centered around 1 µm and nearly all axons clustered within the 1–3 µm diameter range. Quantification of mean axon diameter confirmed these differences. Although the mean axon diameter in the chronic proximal stump (3.80 ± 1.04 μm) was not significantly different from that of uninjured nerves (4.35 ± 0.58 μm), axons in the chronic distal stump were significantly smaller (1.29 ± 0.23 μm) than those in both uninjured nerves (*p* = 0.0042) and chronic proximal stumps (*p* = 0.0110) (Fig. 4B). Myelin thickness was also significantly reduced in the chronic distal stump (0.68 ± 0.07 μm) compared with uninjured nerves (1.60 ± 0.11 μm, *p* = 0.0122) and chronic proximal stumps (1.47 ± 0.38 μm, *p* = 0.0151) (Fig. 4C). Similarly, the g-ratio was significantly lower in the chronic distal stump (0.51 ± 0.10) than in uninjured nerves (0.58 ± 0.03, *p* = 0.0272) but did not differ significantly from the chronic proximal stump (0.54 ± 0.10), indicating a lower ratio of axon diameter to total fiber diameter in the distal compartment (Fig. 4D). Finally, axonal solidity, a measure of axonal shape regularity, was significantly lower in the chronic proximal stump (0.83 ± 0.04) compared with the chronic distal stump (0.91 ± 0.03, *p* = 0.0438), consistent with the irregular and distorted axonal profiles observed histologically in the proximal stump (Fig. 4E). Although axonal solidity was also higher in the chronic distal stump than in uninjured nerves (0.88 ± 0.02), this difference was not statistically significant. Collectively, these findings demonstrate distinct patterns of axonal and myelin remodeling in both compartments following chronic nerve transection, characterized by greater irregularity in nerve fiber morphology in the proximal stump and smaller axon diameters and altered myelin morphometrics in the distal stump.

**Figure 4.**
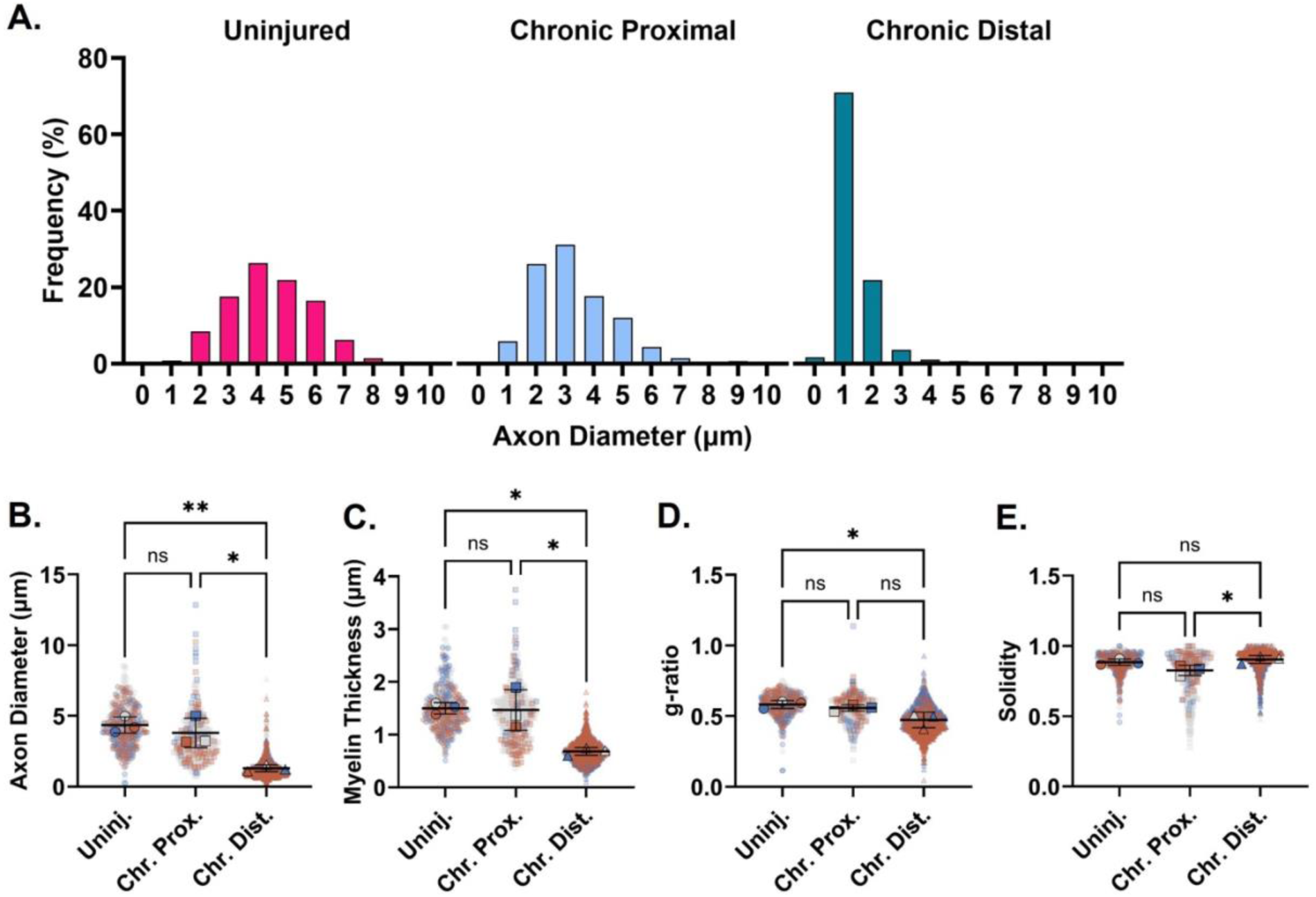
Distinct proximal and distal axon and myelin morphometric profiles. (A) Distribution of axon diameters in uninjured, chronic proximal, and chronic distal nerves. **(B)** Average axon diameter (µm). **(C)** Myelin thickness. **(D)** g-ratio, calculated as axon diameter divided by fiber diameter. **(E)** Axon solidity as a measure of cross-sectional shape regularity. Morphometric measurements were obtained from 100 µm^2^ regions of interest (ROIs) in semithin nerve cross-sections. Four ROIs across three biological replicates were analyzed per condition. Smaller data point are technical replicates; larger data points are biological replicates. Data are presented as mean ± SD. Statistical significance is indicated as * *p* < 0.05.

### The chronic motor nerve transection model recapitulates key features of structural and microenvironmental remodeling in the distal nerve stump

Prolonged nerve discontinuity can produce distinct structural and microenvironmental changes in the proximal and distal stumps; therefore, we next compared these compartments in our chronic injury model. To characterize structural remodeling following chronic facial nerve transection, endoneurial organization and extracellular matrix (ECM) changes were assessed in uninjured nerves and chronic proximal and distal stumps. Laminin immunolabeling highlighted relatively uniform endoneurial tubes in uninjured nerves, while chronic proximal stumps showed predominantly preserved endoneurial architecture with occasional smaller tubes (Fig. 5A). In contrast, chronic distal stumps exhibited widespread endoneurial tube atrophy, with quantitative analysis confirming a significant reduction in average endoneurial tube area (5.67 ± 2.73 μm^2^) compared with both uninjured nerves (25.69 ± 3.30 μm^2^, *p* < 0.0001) and the chronic proximal stump (20.87 ± 4.73 μm^2^, *p* = 0.0007) (Fig. 5B). Another hallmark of chronic denervation, excessive and dysregulated ECM deposition, was examined via collagen I, a primary constituent of endoneurial fibrosis (Vuorinen et al., 1995; Salas-Treviño et al., 2025; Hoang et al., 2026). Chronic distal stumps exhibited excessive and disorganized collagen I deposition within the endoneurium compared with uninjured nerves and chronic proximal stumps (Fig. 5C). Accordingly, quantification revealed a significantly greater collagen I-positive area fraction in the chronic distal stumps (23.55 ± 4.02%) compared with both uninjured nerves (14.10 ± 3.28%, *p* = 0.0116) and chronic proximal stumps (12.69 ± 3.39%, *p* = 0.0051) (Fig. 5D).

**Figure 5.**
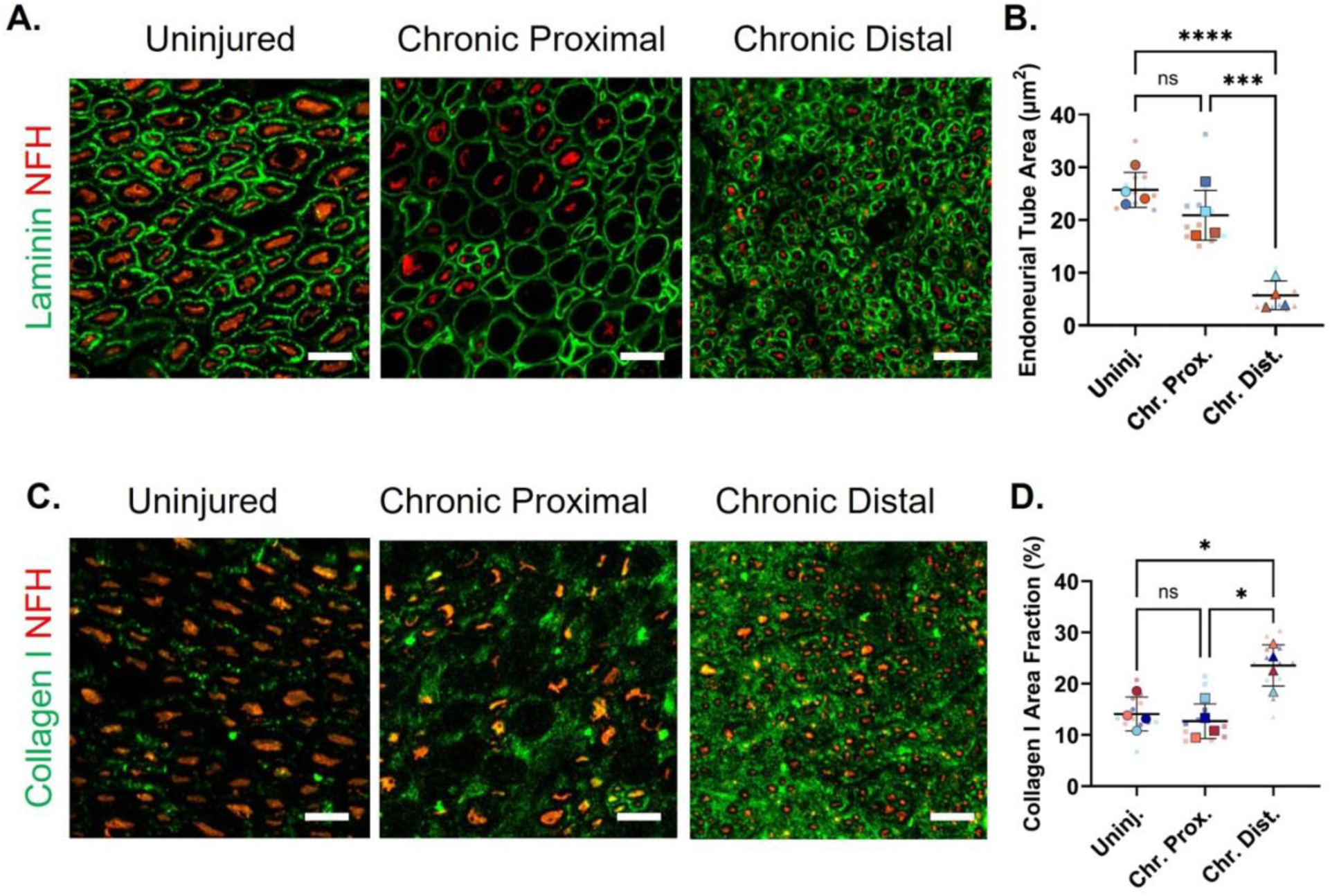
Endoneurial and extracellular matrix remodeling in the chronic distal nerve stumps. Representative immunofluorescence images and corresponding quantification of **(A, B)** laminin (green), highlighting endoneurial tubes and their cross-sectional area (µm^2^), and **(C, D)** collagen I (green), showing extracellular matrix deposition and corresponding area fraction, across Uninjured, Chronic Proximal, and Chronic Distal nerve sections. Laminin and collagen I are co-stained with NFH (red). Scale bar = 10 µm. Data are presented as mean ± SD. Statistical significance is denoted as follows: \**p* < 0.05 and ** *p* < 0.01. NFH, neurofilament heavy chain.

Chronic nerve injury is also associated with vascular remodeling and persistent immune-associated inflammation (Höke et al., 2001; Warner et al., 2026), therefore, we next assessed vascular and immune-cell changes in the proximal and distal stumps of our chronic injury model. Immunofluorescence analysis of PECAM-1, a marker for blood vessels, revealed a non-significant increase in PECAM-1-positive fractional area relative to nerve fascicle area in both the chronic proximal stump (1.43 ± 0.98%) and the chronic distal stump (1.62 ± 1.16%) when compared with uninjured nerves (0.58 ± 0.29%) (Fig. 6A & 6B). The pan-macrophage marker, Iba-1, was then used to label total macrophages (Stratton et al., 2018). Iba-1-positive fractional area was modestly increased in the chronic proximal stump (1.71 ± 0.37%) relative to uninjured nerves (0.87 ± 0.39%), although this difference was not significant, whereas the chronic distal stump (4.81 ± 1.73%) showed a significant increase in Iba-1-positive fractional area compared with uninjured nerves (*p* = 0.0492) (Fig. 6C & 6D). Together, these findings show that our chronic facial nerve transection model exhibits recognized features of chronic nerve remodeling, including distal endoneurial tube atrophy, increased collagen deposition and immune-associated staining, while further demonstrating proximal–distal divergence in the chronic injury microenvironment.

**Figure 6.**
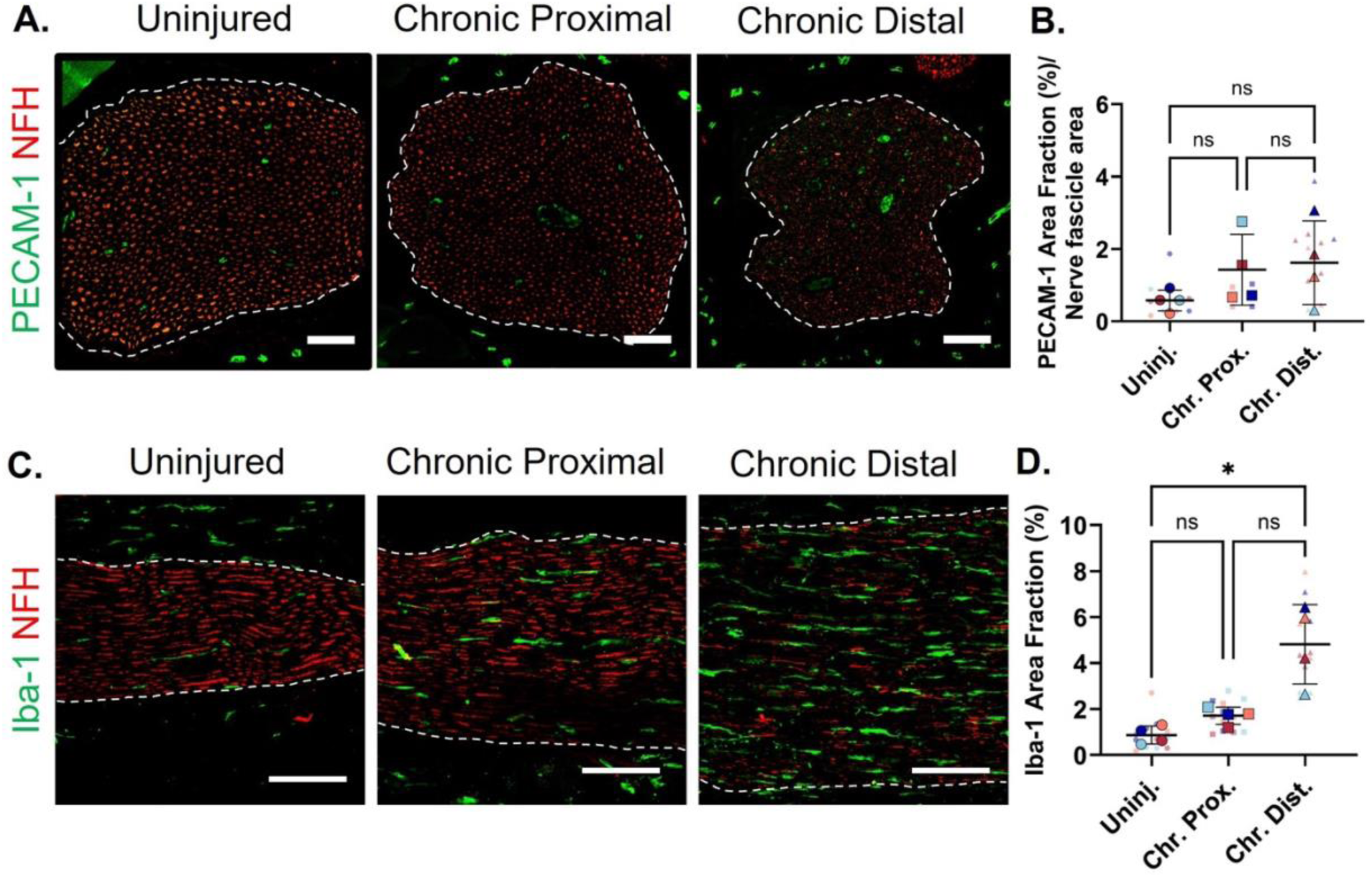
Vascular and inflammatory remodeling in the chronic distal nerve stumps. Representative immunofluorescence images and corresponding quantification of **(A, B)** PECAM-1, highlighting blood vessels and their mean cross-sectional area (µm^2^)/nerve fascicle, and **(C, D)** Iba-1, showing immune cell and corresponding area fraction, across Uninjured, Chronic Proximal, and Chronic Distal nerve sections. PECAM-1 and Iba-1 are co-stained with NFH (red). White dashed lines indicate the boundaries of the nerve tissue. Scale bar = 50 µm. Data are presented as mean ± SD. Statistical significance is denoted as \**p* < 0.05. PECAM-1, platelet endothelial cell adhesion molecule 1; Iba-1, ionized calcium-binding adapter molecule 1; NFH, neurofilament heavy chain.

## Discussion

In this study, we established characterized a rat model of chronic motor nerve transection that recapitulates established pathological hallmarks of prolonged PNI while defining distinct proximal and distal tissue states within the same unrepaired injury. At 10-weeks p.i., the injury model exhibited persistent functional impairment and compartment-specific pathological changes. The proximal stump exhibited terminal traumatic neuroma formation at its terminus, while the more proximal tissue from the neuroma retained NFH expression and relatively preserved nerve architecture. In contrast, the distal stump demonstrated extensive axonal, endoneurial, extracellular matrix, and immune-associated remodeling. Although these pathological features have been described previously, paired characterization of proximal and distal nerve stumps remains limited in chronic motor nerve injury, leaving their distinct structural and microenvironmental responses to prolonged axotomy and denervation incompletely characterized. This gap is compounded by the limited availability of well-characterized, motor-specific chronic injury models that maintain prolonged nerve discontinuity. By establishing such a model and integrating multiparametric analyses of both compartments, this study addresses both gaps, providing a defined experimental framework for investigating chronic motor nerve pathology and evaluating strategies for delayed nerve repair.

Previous experimental studies have established that chronic axotomy and denervation contribute differently to impaired peripheral nerve regeneration. Foundational studies by Fu and Gordon (1995a, 1995b) established the distinct contributions of prolonged axotomy and denervation to impaired peripheral nerve regeneration, while subsequent studies of delayed nerve repair identified compartment-specific alterations in nerve architecture and regenerative support (Sulaiman and Gordon, 2000; Jonsson et al., 2013). Chronic proximal-distal comparisons have also been performed in human nerves (Terenghi et al., 1998) and experimental unrepaired sciatic nerve models (Ruohonen et al., 2005). More recently, Guo *et al*. (2022) characterized the proteome in chronically denervated distal nerves, while Duan *et al*. (2026) demonstrated progressive cellular changes during prolonged denervation. Although these studies have established important aspects of chronic injury biology, they have predominantly utilized mixed-nerve models, heterogeneous clinical samples, or paradigms emphasizing distal regenerative capacity and repair outcomes. The present study extends this literature by systematically characterizing both stumps within a chronic unrepaired motor nerve using complementary structural and microenvironmental assessments.

In our study, the chronic proximal stump exhibited a spatial variation in pathology, consistent with previous findings of extensive axonal disorganization and collagen deposition within terminal neuromas, whereas nerve segments sampled 1 cm proximally retained relatively normal architecture with minimal endoneurial fibrosis (Scott et al., 2022). Similarly, myelinated fiber counts have been shown to progressively normalize with increasing distance from the neuroma, although individual fiber diameters remained smaller even in more proximal regions, indicating that axonal atrophy extends beyond the visibly affected nerve segments (Ko et al., 2011). Consistent with these observations, our chronic proximal stumps exhibited altered axonal caliber, irregular myelin profiles, and reduced axonal solidity relative to distal stumps, despite relatively preserved overall architecture. Furthermore, endoneurial tube area and the area fractions of collagen I, PECAM-1, and Iba-1 did not differ significantly from uninjured nerves. This contrasts with the pronounced fibrotic, vascular, and inflammatory remodeling reported within human neuromas (Deininger et al., 2024), further supporting spatial variation in pathological remodeling along the proximal nerve stump. Taken together, these findings demonstrate that our model recapitulates established features of chronic proximal axotomy, including spatial variation in pathology along the nerve segment.

In contrast, the chronic distal stump exhibited more pronounced axonal and structural remodeling, with substantial reduction in NFH expression and no detectable SCG10, consistent with a chronically denervated distal-stump state marked by progressive axonal degeneration and declining regenerative support (Jonsson et al., 2013; McMorrow et al., 2022; Duan et al., 2026). Despite prolonged separation from the proximal nerve stump, clusters of small-caliber myelinated axons were observed, resulting in a higher myelinated axon density than in uninjured nerves and chronic proximal stumps. Similar residual axonal structures have been reported in chronically denervated distal nerves, including occasional sprouts despite experimentally restricted regeneration and thinly myelinated axons in chronically transected human nerves, although these findings are variable and do not by themselves indicate effective regeneration (Röyttä and Salonen, 1988; Terenghi et al., 1998). Additionally, chronic distal stump fibers exhibited reduced myelin thickness alongside lower g-ratios, likely reflecting disproportionately smaller axon diameters rather than increased absolute myelin thickness. Collateral sprouts from neighboring sensory and autonomic nerve pathways may explain the small-caliber myelinated axons observed. Previous studies have demonstrated that intact sensory axons can undergo collateral sprouting into denervated territories (Lemaitre and Court, 2021; Jeon et al., 2024). Additionally, parasympathetic fibers traveling via the sensory infraorbital nerve can reinnervate denervated facial muscles in rats (Tereshenko et al., 2022). Together, these findings suggest potential non-motor sources of axonal ingrowth, although the precise origin of the observed fibers remains uncertain and warrants further investigation through retrograde tracing.

Beyond axonal changes, the chronic distal stump exhibited substantial endoneurial and microenvironmental remodeling. At 10 weeks p.i., endoneurial tubes were markedly smaller but retained recognizable laminin-defined boundaries, indicating atrophy with preservation of the underlying architecture. This is consistent with previous reports of Schwann cell column shrinkage and persistent endoneurial remnants during chronic denervation (Röyttä and Salonen, 1988; Vuorinen et al., 1995). Increased collagen I deposition further indicated extracellular matrix remodeling, consistent with previous histological and proteomic findings (Jonsson et al., 2013; Guo et al., 2022). Increased Iba-1-positive labeling was also observed, in line with reported immune-associated changes following nerve injury and during chronic denervation (Ruohonen et al., 2005; Zhao et al., 2022; Ouyang et al., 2025), although this alone does not establish immune cell activation or state. Collectively, these findings demonstrate that our model recapitulates established hallmarks of chronic distal nerve pathology, including endoneurial atrophy, extracellular matrix remodeling, and immune-associated alterations.

A major strength of this study is the use of the rat facial nerve buccal branch as a motor-specific model of chronic PNI, offering several experimental advantages over commonly used mixed sensory–motor nerves, such as the sciatic nerve. Its predominantly motor composition minimizes sensory-related confounding effects, while the reduced risk of autotomy improves tolerability for longitudinal studies (Merkulyeva, 2026), Furthermore, whisker movement provides a direct, noninvasive functional readout of facial motor impairment that can be assessed longitudinally (Olmstead et al., 2015; Bengur et al., 2022). The facial nerve also offers practical anatomical advantages, as its surgical accessibility, relatively fixed position, and reduced susceptibility to mechanical disruption compared with limb nerves facilitate consistent surgical manipulation, tissue sampling, and experimental interventions (Olmstead et al., 2015; Ali et al., 2020; Petrov et al., 2022). Another key strength is the clinical relevance of the facial nerve model to facial paralysis and facial nerve reconstruction (Xie et al., 2024). The 10-week chronic injury endpoint also captures prolonged denervation and associated pathological changes seen in human nerve injuries, which may be underrepresented in acute rodent repair models due to shorter regenerative distances and faster axonal growth rates (Marsh et al., 2024). Additionally, sampling nerve segments away from the terminal transection ends more closely approximates the tissue potentially retained following surgical debridement during nerve reconstruction (Fones et al., 2024). Finally, the paired, multiparametric characterization of proximal and distal nerve stumps at a chronic timepoint represents a key strength of this study. By integrating structural and microenvironmental analyses across both compartments within the same experimental model, this approach builds upon existing literature and provides a more complete characterization of chronic motor nerve pathology, considering the pathological consequences of prolonged axotomy and denervation together rather than in isolation.

A few limitations of this study are worth considering when interpreting the findings. First, analyses were restricted to a single chronic endpoint at 10 weeks p.i., limiting assessment of the temporal progression of proximal and distal pathology. Chronic denervation is a dynamic process involving progressive remodeling of the distal nerve microenvironment, declining Schwann cell regenerative capacity, and cellular deterioration (Xu et al., 2023; Duan et al., 2026). Thus, the pathological features observed at 10 weeks p.i. may differ from those at earlier or later chronic stages. Second, tissue sampling was restricted to defined regions away from the terminal transection ends, limiting assessment of spatial variation in pathological remodeling along the longitudinal axis of the peripheral nerve within either the proximal or distal stump. This limitation is particularly relevant given previous studies demonstrating that cellular responses, molecular signaling, and tissue remodeling can vary with distance from the injury site, with distinct changes occurring near the cut surface compared with more distant nerve segments (Sheu et al., 2000; Spezia et al., 2026). Third, our functional assessment addressed a major source of overlapping whisker-pad innervation by transecting the marginal mandibular branch (Marin et al., 2026). Nevertheless, whisker movement may still be influenced by other nerve pathways and compensatory responses and therefore cannot be attributed exclusively to buccal branch function (Heaton et al., 2014; Rink-Notzon et al., 2022). Accordingly, future studies using this buccal branch-specific model to assess functional recovery should consider potential contributions from other nerve sources and evaluate whisker movement alongside evidence of axonal regeneration through the repaired buccal branch and reinnervation of the vibrissae musculature. Finally, rodent models do not fully recapitulate human nerve injury, because human nerves must regenerate across much longer distances and under less permissive chronic denervation conditions (Scheib and Höke, 2013). Nevertheless, the use of a chronic injury endpoint more closely approximates the prolonged denervation-associated pathological changes observed clinically, which are often underrepresented in acute rodent models (Marsh et al., 2024).

Overall, this model reproducibly generated distinct chronic proximal and distal tissue states, providing an experimental platform to define how chronic nerve injury evolves over time and influences later regenerative outcomes. Future studies should incorporate additional chronic timepoints and spatially resolved sampling at defined distances from the injury site to determine when and where pathological remodeling emerges and how the local regenerative microenvironment changes. A key unresolved question is which chronic tissue changes, beyond the established loss of Schwann cell regenerative support (Fuentes-Flores et al., 2023), are merely markers of denervation and which causally drive regenerative failure and define actionable therapeutic windows. Addressing this question will require pairing spatiotemporal characterization with delayed-repair experiments and targeted interventions, for example, strategies to preserve or restore Schwann cell regenerative support (McMorrow et al., 2022; Rebolledo et al., 2026), to test whether specific chronic tissue changes causally limit regenerative outcomes. Ultimately, defining the spatial and temporal features of chronic nerve injury that constrain regeneration could identify therapeutic windows and compartment-specific targets that inform future pharmacologic strategies and bioactive nerve guidance conduit design.

## Conclusion

In conclusion, this study establishes and characterizes a chronic unrepaired motor nerve transection model, enabling direct comparison of the proximal and distal nerve stumps following prolonged discontinuity. Through multiparametric analyses, the model demonstrates distinct patterns of axonal remodeling, myelin pathology, and microenvironmental alterations across both compartments, recapitulating established features of chronic nerve injury. These findings provide a foundation for future studies to investigate the temporal progression of proximal and distal pathology, identify the cellular and molecular mechanisms underlying chronic remodeling, and determine how these changes influence regenerative outcomes following delayed nerve repair. Furthermore, the surgical accessibility and tolerability of the facial nerve model support its use in evaluating therapeutic strategies targeting the proximal stump, distal stump, or both. Collectively, these findings reinforce the importance of considering chronic nerve injury as a two-compartment process characterized by distinct but interrelated pathological changes.

## Acknowledgments

This research was generously supported by the Stanford Department of Otolaryngology - Head & Neck Surgery and philanthropic contributions from Daniel and Diane Riccio, to whom the authors are deeply grateful.

## Author contributions

**Conceptualization:** C.Y., and J-P.P. **Investigation:** C.Y., and A.N. **Methodology:** C.Y. **Formal Analysis:** C.Y., and L.J. **Visualization:** C.Y., and L.J. **Writing – Original Draft Preparation:** C.Y., and A.N. **Writing – Review & Editing:** C.Y., and J-P.P. **Funding Acquisition, Supervision, and Resources:** J-P.P. All authors approved the final version of this manuscript.

## Conflicts of interest

The authors declare no competing interests.

## Declaration of AI and AI-assisted technologies in the writing process

During the preparation of this manuscript, the authors used ChatGPT (GPT-5.6 Sol; OpenAI) to assist with grammar checking. The authors subsequently reviewed and edited the text as necessary and take full responsibility for the content of the publication.

## Data availability statement

All relevant data are within the paper.

---

**Supplementary Figure 1.**
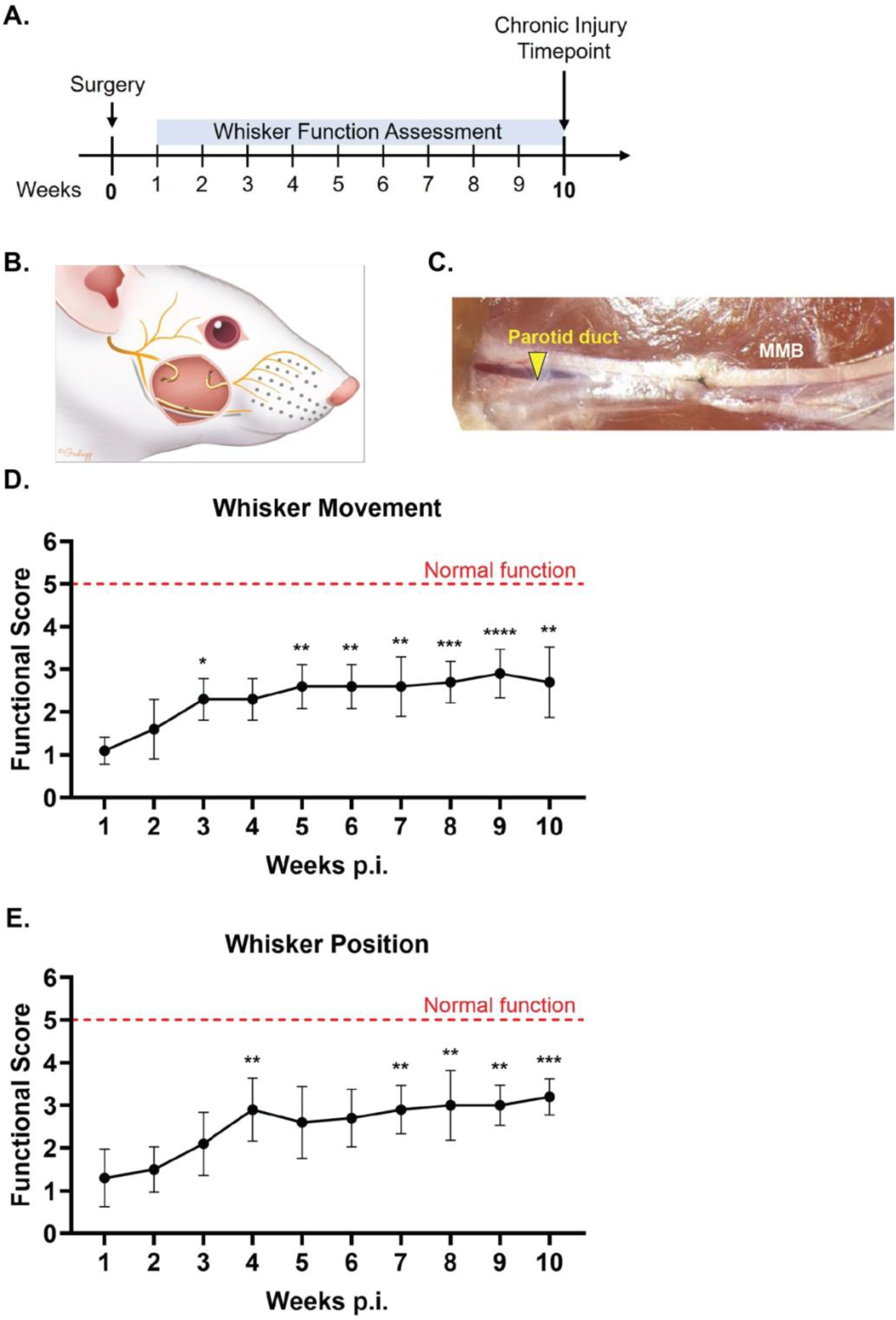
Persistent whisker dysfunction is observed in the chronic facial nerve transection model. **(A)** Schematic of the experimental timeline for whisker function analysis. Rats were video-recorded weekly from 1- to 10-weeks p.i. to assess functional recovery. **(B)** Complete transection of the buccal branch, followed by diversion and suture of the proximal and distal nerve ends. The marginal mandibular branch (MMB) was also transected and repaired to temporarily eliminate additional motor input to the vibrissae musculature. **(C)** Surgical manipulation of the MMB may be complicated by its close association with the parotid duct. The duct is not readily visible intraoperatively but can be identified postmortem. **(D)** Whisker movement scores over the 10-week post-injury period. **(E)** Whisker position scores over the 10-week post-injury period. Whisker function was scored on a 1–5 scale, with 1 indicating no function and 5 indicating complete function. Data are presented as mean ± SD. A cohort of ten rats (n = 5 males and n = 5 females) was used for functional analysis. Functional scores were analyzed using a Friedman test followed by Dunn’s multiple-comparisons test, with each time point compared with 1-week p.i. Statistical significance is denoted as follows: * *p* < 0.05, \*\**p* < 0.01, *** *p* < 0.001, and **** *p* < 0.0001. BB, buccal branch; MMB, marginal mandibular branch; p.i., post-injury. Rat illustration (B) published with permission from Ms. Christine Gralapp.

## Notes

### Competing Interest Statement

The authors have declared no competing interest.

